# Retromer repletion with AAV9-VPS35 restores endosomal function in the mouse hippocampus

**DOI:** 10.1101/618496

**Authors:** Yasir H. Qureshi, Diego E. Berman, Ronald L. Klein, Vivek M. Patel, Sabrina Simoes, Suvarnambiga Kannan, Rebecca Cox, Samuel D Waksal, Beth Stevens, Gregory A. Petsko, Scott A. Small

**Affiliations:** Departments of Neurology and the Taub Institute for Research on Alzheimer’s Disease and the Aging Brain, Columbia University; Department of Pharmacology, Toxicology and Neuroscience, Louisiana State University Health Sciences Center; Department of Neurology and Neuroscience, Weill Cornell Medical School, Cornell University; Meira GTX, Alexandria Center for Life Sciences, New York; Department of Neurology, Harvard Medical School and the Howard Hughes Medical Institute, Harvard University

## Abstract

Retromer has emerged as a master conductor of endosomal trafficking, and VPS35 and other retromer-related proteins are found to be deficient in late-onset Alzheimer’s disease (AD). Depleting VPS35 in neurons impairs retromer function, affecting for example the trafficking of the amyloid-precursor protein (APP) and the glutamate receptor GluA1. Whether VPS35 repletion, after chronic *in vivo* depletion, can rescue these impairments remains unknown. Here we set out to address this question by using a viral vector approach for VPS35 repletion. First, we completed a series of studies using neuronal cultures in order to optimize AAV9-VPS35 delivery, and to understand how exogenous VPS35 expression affects its endogenous levels as well as its binding to other retromer proteins. Next, we completed a series of studies in wildtype mice to determine the optimum protocol for *in vivo* delivery of AAV9-VPS35 to the hippocampus. We relied on this information to deliver AAV9-VPS35 to the hippocampus of mice genetically engineered to have chronic, neuronal-selective, VPS35 depletion. VPS35 repletion in the hippocampus was found to normalize APP cleavage and to restore glutamate receptor levels. Unexpectedly, chronic VPS35 depletion in neurons caused glial activation, similar to the pattern observed in AD, which was also partially normalized by VPS35 repletion. Taken together, these studies strengthen the mechanistic link between retromer and AD, and have therapeutic implications.

## INTRODUCTION

Retromer is a multi-modular protein assembly that recycles transmembrane proteins out of the endosomal compartment^1^. The ‘cargo recognition core’ is a central module of retromer function because it is the core to which transported cargo binds, but also because it is with this core that other retromer modules interact^2^. This heterotrimeric core includes the protein ‘Vacuolar Protein Sorting 35’ (VPS35) to which the other core proteins VPS29 and VPS26 bind. Depleting VPS35 causes a secondary reduction in the other core proteins and impairs retromer function^1^.

VPS35 and other retromer core proteins have been found deficient in the hippocampal formation of patients with late-onset Alzheimer’s disease (AD)^3^. When acutely modeled in neurons and cell culture, retromer depletion increases the residence time of the amyloid-precursor protein (APP) in endosomes, accelerating its proteolytic cleavage^4–7^. Other studies have depleted neuronal VPS35 in hippocampal slices to show that retromer is required for the recycling of the glutamate receptor, ionotropic, AMPA1 alpha 1 (GluA1, also called GluR1) to the cell surface^8^. Since reducing receptor recycling accelerates its degradation, retromer deficiency might account for the GluA1 deficiency observed in AD, a defect that is thought to mediate synaptic pathology^9–12^ in AD.

These and other studies support the conclusion that the chronic VPS35 deficiency observed in AD impairs retromer function and can contribute to the neuronal defects observed in the disease. Lacking, however, is evidence showing that VPS35 repletion, after chronic *in vivo* depletion, restores retromer function. We therefore generated two new experimental tools: A novel mouse model in which VPS35 is selectively depleted in neurons, and an AAV9 vector that carries the human VPS35 plasmid with which to replete VPS35.

Before beginning *in vivo* repletion studies, however, we first needed to complete an extensive series of studies in neuronal culture and wildtype mice to optimize the VPS35 transduction protocol, and to understand its consequences in non-depleted neurons. We then completed *in vivo* VPS35 repletion studies, and find that it restores retromer function, normalizing APP cleavage and glutamate receptor levels in the mouse hippocampus. These studies also unexpectedly showed that VPS35 depletion in neurons secondarily activated glia, similar to the pattern observed in AD, and that repletion of VPS35 in neurons partially abrogated this glial response.

## RESULTS

### AAV9-VPS35 expression and the endogenous retromer

To determine expression efficacy and optimum delivery, we first C-terminally tagged human VPS35 with green fluorescent protein (GFP), and delivered AAV9-VPS35-GFP to mouse neuronal cultures. Immunoblot levels of endogenous and exogenous VPS35 were measured across three doses. A pattern consistent with autoregulation was observed (Fig. 1A), whereby the higher the levels of *exogenous* VPS35 expressed, the lower the levels of *endogenous* VPS35 were detected. A significant increase in total VPS35 was observed at both the middle dose (t=10.74, p=0.0004) and highest dose (t=26.02, p=0.00001).

**Figure 1.**
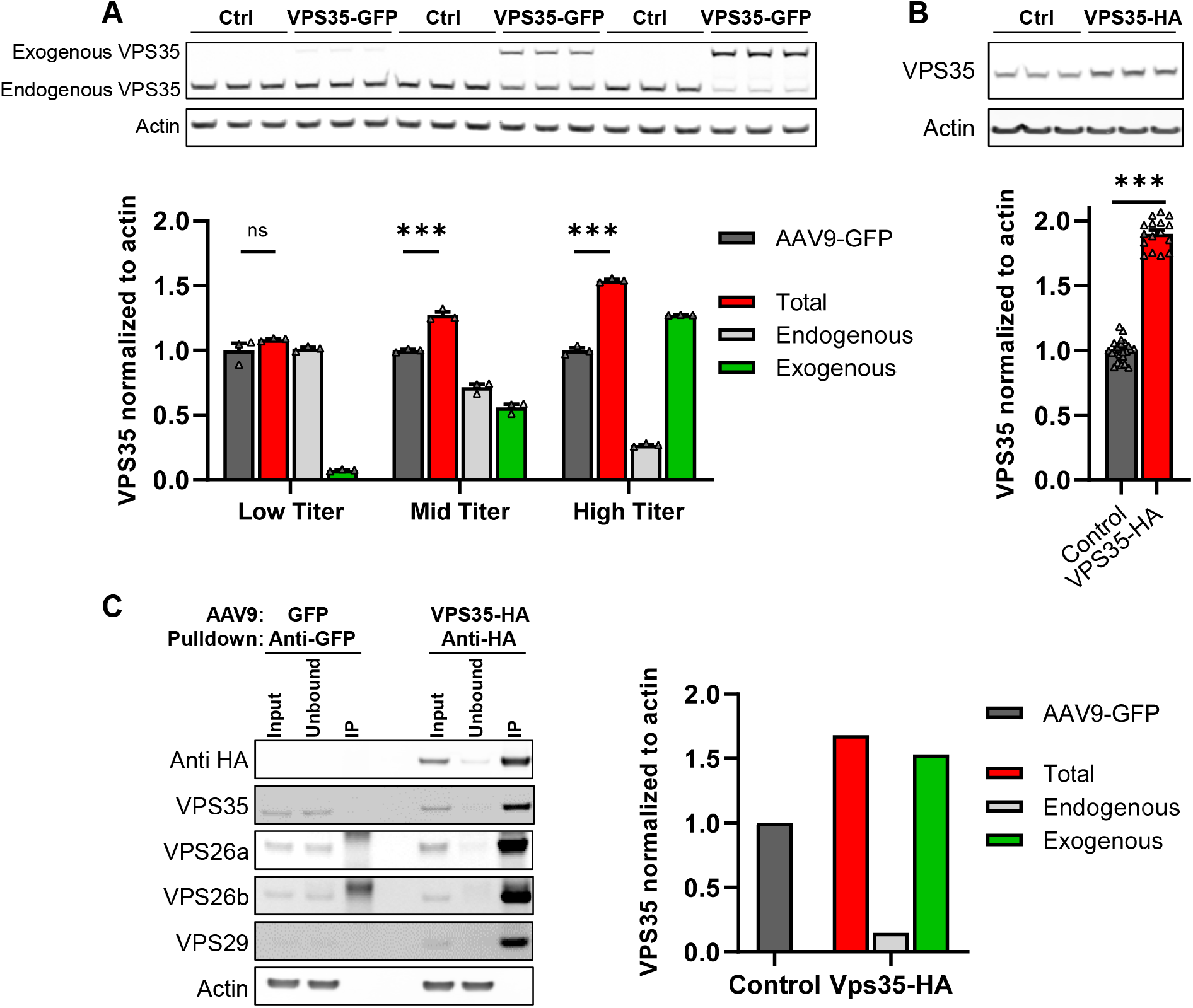
AAV9-VPS35 expression and the endogenous retromer. **(A)** Immunoblots showing expression levels of exogenous and endogenous VPS35 after delivery of 3 doses of AAV9-VPS35-GFP in primary neuronal culture (5E+9 vg/well, 2E+10 vg/well, 1E+11 vg/well), and after delivery of AAV9-GFP as a control. Bar graphs show the mean levels of endogenous VPS35 (grey bars), exogenous VPS35 (green bars) and total VPS35 (red bars). Mean level of VPS35 with AAV9-GFP expression (dark grey bars) shown as control. **(B)** Representative immunoblots showing expression levels AAV9-VPS35-HA at 1E+11 vg/well. Bar graphs show mean levels, normalized by actin and AAV9-GFP and AAV9-EV (empty vector) were used as controls **(C)** Immunoblots showing that anti-HA antibody co-immunoprecipitated endogenous retromer core proteins--VPS26a, VPS26b, and VPS29. Bar graphs represent levels of VPS35, total (red; input), endogenous VPS35 (light gray; unbound), and exogenous VPS35 (green; unbound subtracted from input) extracted from the co-immunoprecipitation data. Control bar (dark gray; input from the control reaction)

Because VPS35 is a protein that acts as a platform to which other retromer proteins bind, we next tagged the human VPS35 with HA, since it is smaller than GFP and less likely to interfere with VPS35’s function. This size difference has also been reported to influence expression levels^13–17^. We find that AAV9-VPS35-HA, delivered at similar doses, caused higher total VPS35 expression than AAV9-VPS35-GFP (Fig. 1B). A co-immunoprecipitation study was performed, using anti-HA antibody, showing that exogenously expressed VPS35 pulls down endogenous VPS29 and VPS26 (Fig. 1C), thus establishing that exogenous HA-tagged VPS35 binds endogenous retromer core proteins.

### AAV9-VPS35 delivered to the mouse hippocampus

Next, to determine the optimum *in vivo* AAV9-VPS35 delivery we completed 8 experiments (3-6 animals per experiment), delivering AAV9-VPS35-HA intracranially to the dorsal CA1 region of the hippocampus of wildtype mice in varying doses and volumes. Brain slices were generated for fluorescent immunohistochemistry (IHC) and probed with anti-HA antibody to map the degree and distribution of expression, using neuronal and nuclear markers to detect potential toxicity. In parallel, tissue samples from the CA1 were microdissected and processed for immunoblotting to quantify expression levels (Fig. 2A - Right).

**Figure 2.**
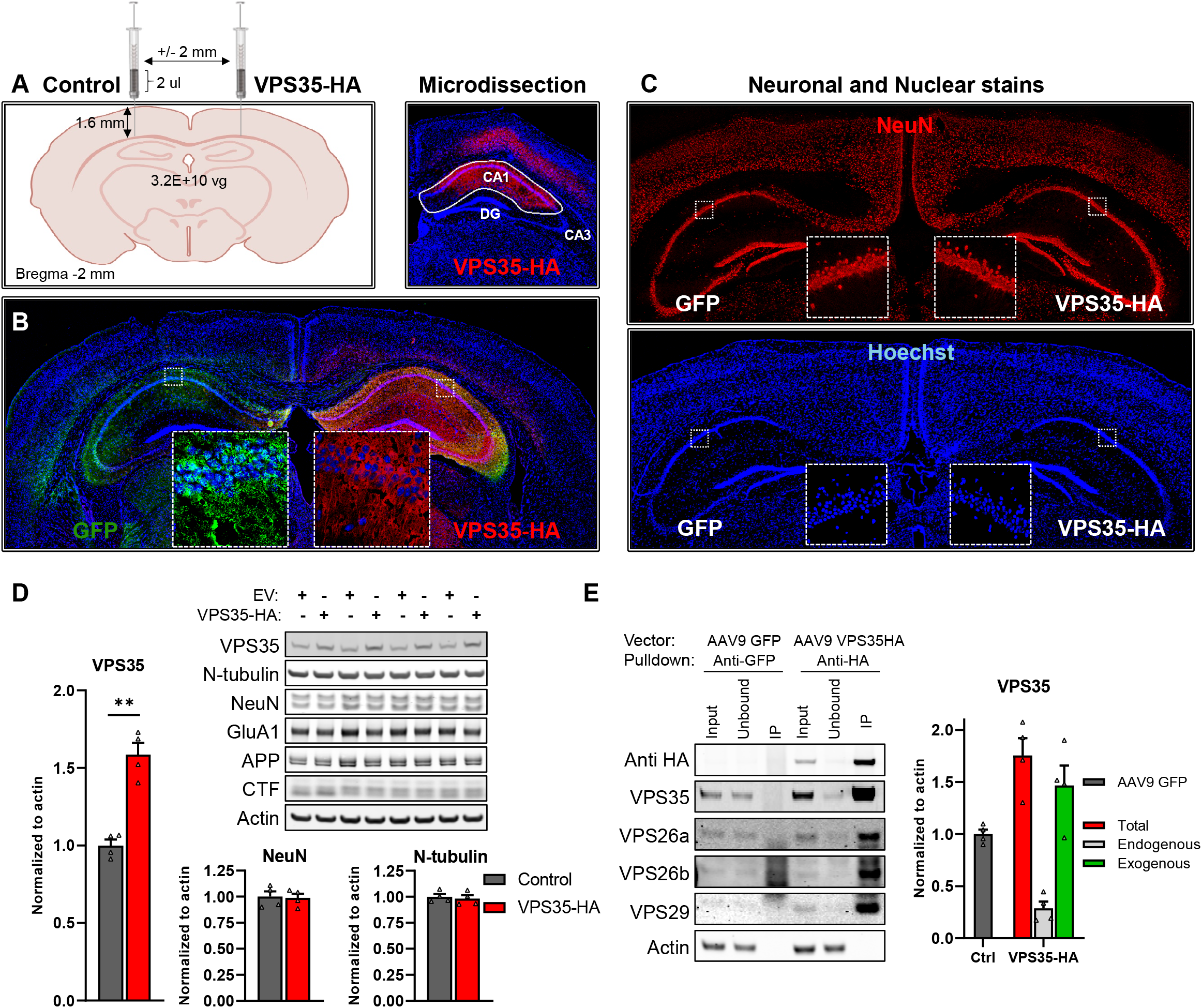
AAV9-VPS35 delivered to the mouse hippocampus. **(A)** Schematic representation (left panel) of the protocol of AAV9 delivery into the hippocampal CA1 region. All animals were injected at 3 months, aged to 6 months of age, at which time brains were processed for fluorescent IHC and biochemistry (as illustrated in the right panel). **(B)** Coronal brain sections immunofluorescently stained 3 months after AAV9-VPS35-HA into right CA1 and AAV9-GFP into the left CA1, showing expression distribution of exogenous VPS35. Insets represent higher magnification images of indicated subregions **(C)** Brain sections stained with the neuronal marker, NeuN and the nuclear marker, hoechst, showed no changes in the neuronal layers compared to the contralateral CA1. **(D)** Immunoblots showing VPS35 overexpression and levels of beta III tubulin (N-tubulin), NeuN, glutamate receptor ionotropic AMPA1 alpha 1 (GluA1), full length APP (APP), C-terminal fragments of APP (CTF), and Actin in the CA1 regions injected with AAV9 empty vector (EV) or VPS35-HA. **(E)** Immunoblots showing that anti-HA antibody co-immunoprecipitated endogenous retromer core proteins -- VPS26a, VPS26b, and VPS29. Bar graphs represent levels of VPS35, total (red=input), endogenous VPS35 (light grey=unbound), and exogenous VPS35 (green = unbound subtracted from input) extracted from the co-immunoprecipitation data. Control bar (dark grey=input from the control reaction).

Using the optimum delivery protocol (Fig. 2A, left panel), we injected AAV9-VPS35-HA to 3-month old mice that were then aged to 6 months. Results showed a broad expression distribution across the dorsal hippocampus, with a significant increase in VPS35 (t=8.74, p=0.003) with no evidence of toxicity (Fig. 2B, C&D). As in neuronal culture, a co-immunoprecipitation study with anti-HA antibody showed that exogenous VPS35 downregulates the corresponding endogenous protein, and that exogenous VPS35 binds other retromer core proteins (Fig. 2E).

### VPS35 depletion-repletion and neuronal phenotypes

By crossing mice expressing loxP-flanked VPS35 (*VPS35*^fl/fl^) with mice expressing *Cre* recombinase under the *Camk2α* promoter we generated VPS35 neuronal-selective knockout (VPS35 nsKO) mice (Suppl. Fig.), in whom VPS35 is depleted in forebrain neurons including the hippocampus (Fig. 3A). Using the optimized protocol, AAV9-VPS35-HA was delivered to the right CA1 region of the VPS35 nsKO mice, and AAV9-GFP was delivered to the contralateral CA1 as control (Fig. 3D). In addition, *VPS35*^fl/fl^ mice were evaluated as non-injected controls.

**Figure 3.**
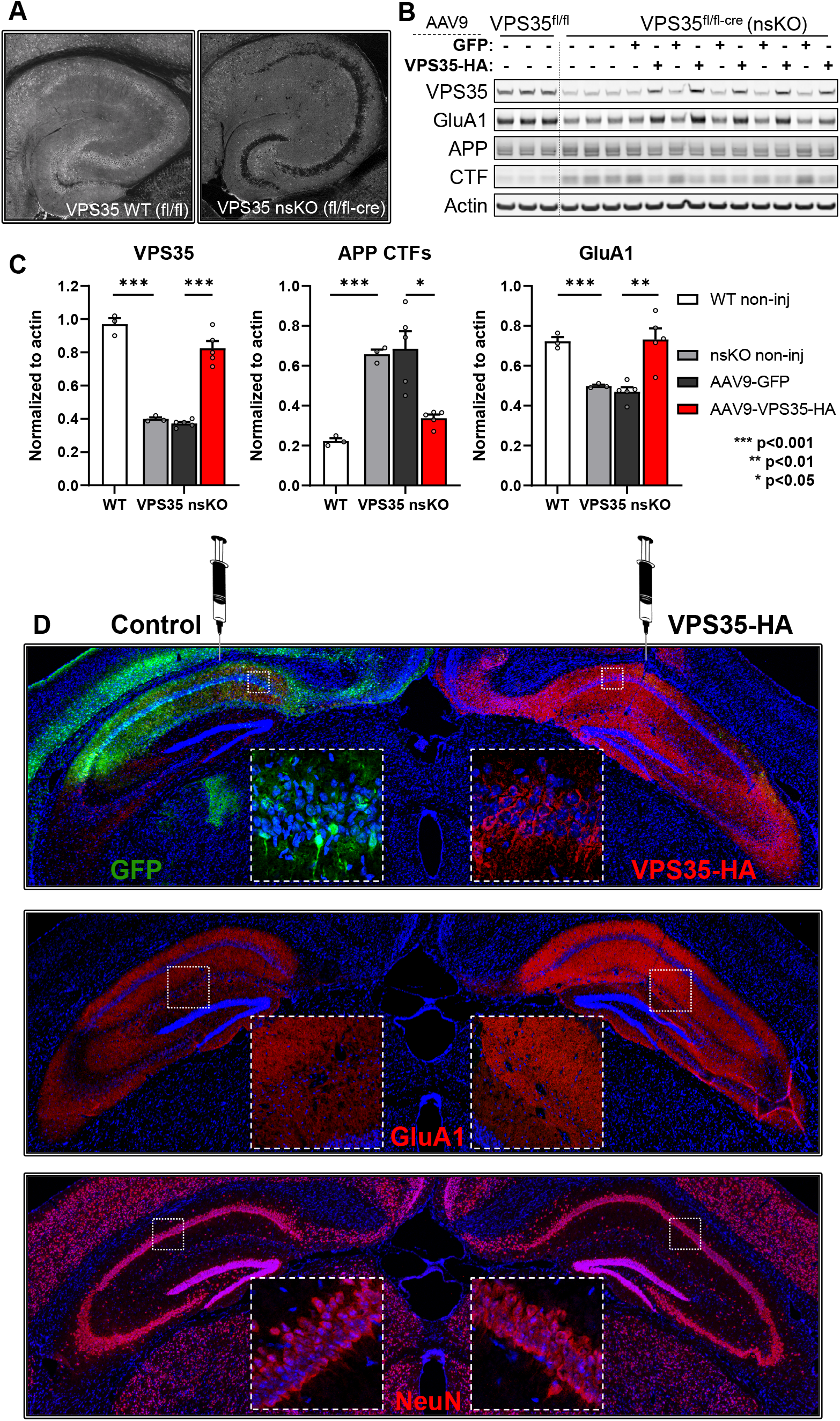
Neuronal phenotypes in VPS35 nsKO mice. **(A)** Hippocampal section from WT and VPS35 neuronal-selective knockout (VPS35 nsKO) mice stained with anti-VPS35 antibody show a complete depletion of VPS35 in the neuronal layers. **(B)** Immunoblots from of the CA1 region of WT (VPS35^fl/fl^ n=3) and VPS35 nsKO (n=8) mice (6 months of age) showing VPS35, GluA1, full length APP, C-terminal fragments of APP (CTF) and Actin protein levels. AAV9-GFP and AAV9-VPS35-HA were injected in left and right CA1 respectively in 5 of the VPS35 nsKO mice at 3 months. **(C)** Bar graphs represent actin-normalized levels of VPS35 (left graph), C-terminal fragments of APP (APP CTFs; middle graph), and glutamate receptor ionotropic AMPA1 (GluA1; right graph). WT=VPS35^fl/fl^. **(D)** Coronal brain sections immunofluorescently stained with anti-HA and anti-GFP, 3 months after AAV9-VPS35-HA into right CA1 and AAV9-GFP into the left CA1 of VPS35 nsKO mouse (upper panel), showing increase in GluA1 expression in the right hippocampus (middle panel), with no detectable changes in neuronal staining (lower panel)

All mice were injected at 3 months of age and aged to 6 months. VPS35 depletion accelerated APP cleavage, indicated by an increase in its C-terminal fragments (CTFs) (t=15.89, p=0.00009), and a striking reduction in GluA1 levels (t=9.70, p=0.0006). VPS35 repletion for 3 months with AAV9-VPS35-HA normalized APP misprocessing (t=3.88, p=0.018), and completely restored GluA1 levels as shown by immunoblotting (t=6.45, p=0.003), (Fig. 3B&C), and confirmed by fluorescent IHC (Fig. 3D).

### VPS35 depletion-repletion and glia responses

Depleting neuronal VPS35 was observed to cause an activation of astrocytes, as detected by an increase in GFAP (t=2.74, p=0.052) (Fig. 4A). VPS35 repletion normalized this response, shown by immunoblotting (t=3.39, p=0.028) and fluorescent IHC (Fig. 4A,C&D). Exogenous VPS35-HA expression in WT mice had minimal effect on astrocytes at 3 months post-surgery (Fig. 4B).

**Figure 4.**
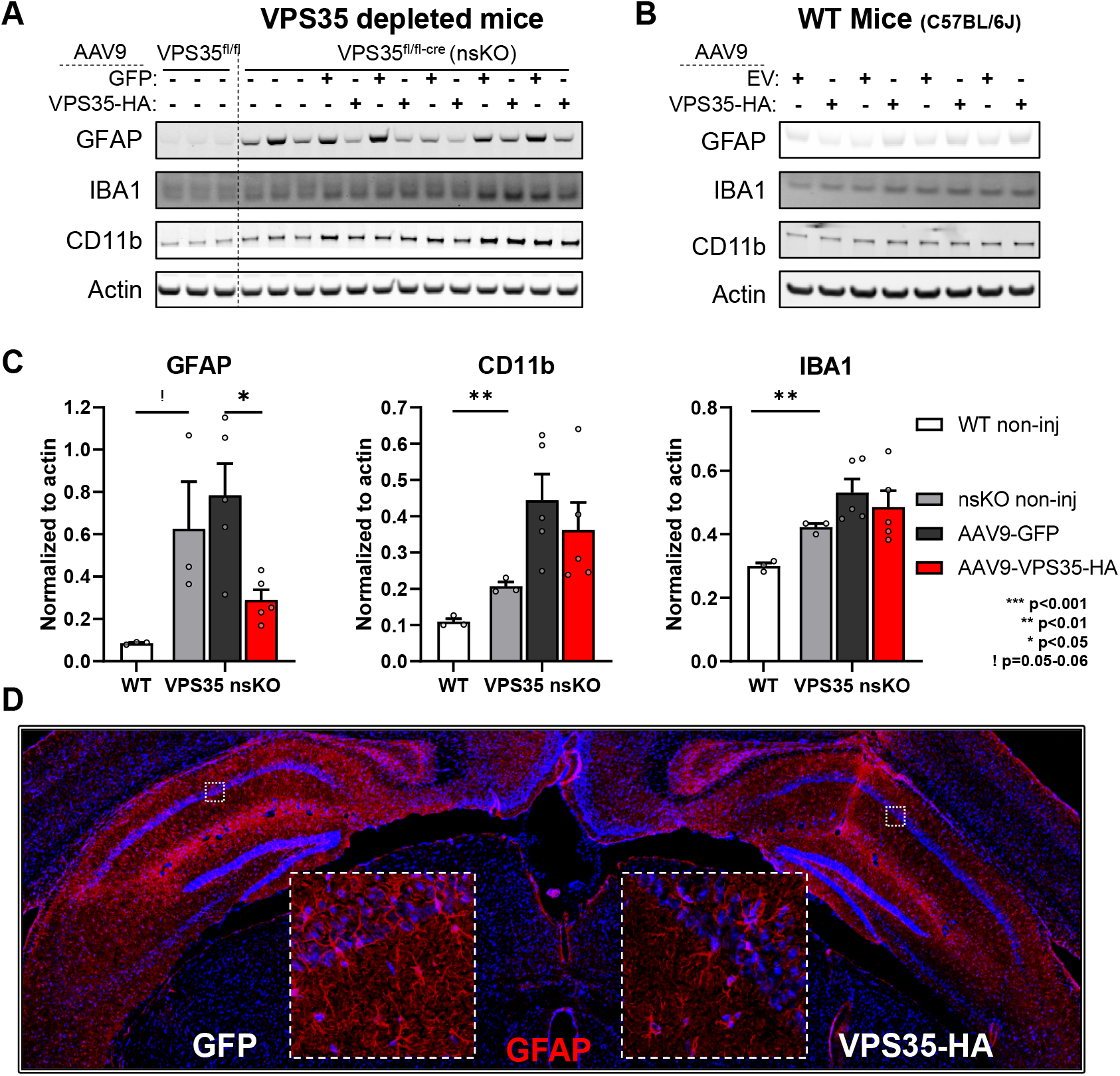
Glial phenotypes in VPS35 nsKO mice. **(A)** Immunoblots from the CA1 region of WT (VPS35^fl/fl^ n=3) and VPS35 nsKO (n=8) mice (6 months of age) were probed for astrocyte marker glial fibrillary acidic protein (GFAP), and microglial markers Ionized calcium-binding adapter molecule 1 (IBA1), and CD11 antigen-like family member B (CD11b). AAV9-GFP and AAV9-VPS35-HA were injected in left and right CA1 respectively in 5 of the VPS35 nsKO mice at 3 months. VPS35 expression in these animals is shown in Fig. 3B **(B)** Immunoblots from the CA1 region (WT=C57BL/6J) injected with AAV9 empty vector (EV) or VPS35-HA probed for astrocyte and microglial markers. VPS35 expression in these animals is shown in Fig. 2D. **(C)** Bar graphs represent actin-normalized levels of the three glial markers from panel A. VPS35 nsKO mice displayed a dramatic increase in gliosis, which was partially normalized by VPS35 repletion (WT=VPS35^fl/fl^). **(D)** Coronal brain section immunofluorescently stained, 3 months after AAV9-VPS35-HA injection into right CA1 and AAV9-GFP into the left CA1 of VPS35 nsKO mouse showing a decrease in GFAP expression in the right hippocampus, confirming the lower astrocytic signal detected in immunoblots with VPS35-HA overexpression.

Depleting neuronal VPS35 also caused an activation of microglia, as detected by an increase in IBA1 (t=8.24, p=0.001) and CD11b (t=6.65, p=0.002) levels (Fig. 4A&C). Immunoblotting showed that VPS35 repletion had a modest, but not statistically significant, effect on IBA1 and CD11b. Again, exogenous VPS35-HA expression in WT mice had minimal effects on microglia at 3 months post-surgery (Fig. 4B)

By co-labelling we confirmed previous studies, showing that when AAV9 is injected directly into the brain parenchyma it is not expressed in microglia and minimally expressed in astrocytes^18–22^ (Fig. 5A&B).

**Figure 5.**
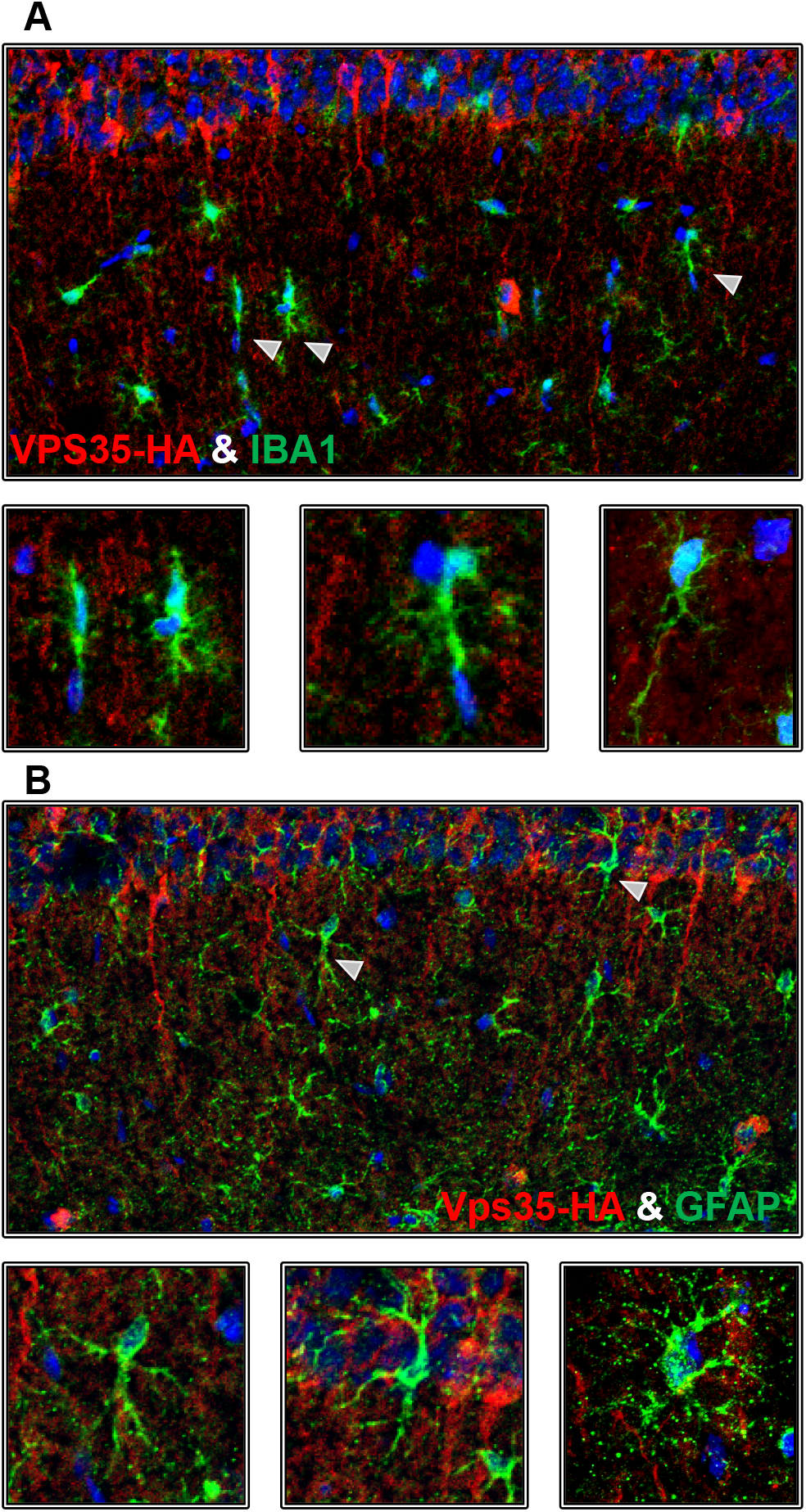
AAV9-VPS35-HA cell specificity. Coronal brain sections immunofluorescently stained, 3 months after AAV9-VPS35-HA injection into right CA1 of VPS35 nsKO mouse showing a lack of co-localization of VPS35-HA with microglial marker IBA1 **(A)**, and minimal overlap with astrocyte marker GFAP **(B)**.

## DISCUSSION

A range of previous studies have depleted VPS35 to model the retromer deficient state observed in the hippocampal formation of patients with late-onset AD^1^. Lacking, however, have been studies that show that *in vivo* VPS35 repletion restores retromer function in neurons, and corrects AD-associated defects. The generation of a novel mouse model in which VPS35 is chronically and selectively depleted in mice forebrain neurons provide this missing evidence.

A recent study showed that viral vector delivery of VPS35 to a neonatal triple-transgenic mouse model rescued defects induced by amyloid and tau pathology^23^. These genetically complex mice overexpress two human mutations in APP and a mutation in presenilin1 (PSEN1), each a cause of autosomal-dominant AD; and, overexpresses a human mutation in the microtubule associated protein tau (MAPT) that is a cause of fronto-temporal degeneration (FTD). Despite concerns raised about non-physiological overexpression of multiple mutations in three different genes linked to AD and to FTD, this mouse has proven extremely useful to model the consequences of aggressive amyloid and tau pathology in the context of autosomal-dominant disease^24–26^. Our findings compliment this prior study^23^, by not only investigating a mouse that is genetically simpler -- depleting a single endogenous gene -- but by using a model that better reflects ‘sporadic’ late-onset AD.

By documenting APP misprocessing and GluA1 deficiency in the hippocampus of neuronal-selective VPS35 depleted mice, and more importantly, by rescuing these defects with VPS35 repletion, our findings strengthen the causal conclusion that retromer deficiency in late-onset AD can mediate these neuronal phenotypes of the disease. Unexpectedly, we also provide evidence for a trans-cellular effect, whereby defects in the neuronal retromer can mediate astrocyte and microglia activation, a glial phenotype that is characteristic of AD. We note that while VPS35 depletion caused an activation of both astrocytes and microglia, the abrogating effect of VPS35 repletion was observed more reliably for astrocytes. Whether an effect on microglia would require earlier VPS35 repletion, or that once induced a shift in microglial state is less sensitive to VPS35 repletion, remains unknown. The mechanisms by which a depletion in neuronal retromer mediates the glial response, and whether this trans-cellular effect depends on the neuronal retromer in particular, is also unknown. However, since human genetics have established that glia can modulate AD risk^27^, our studies show how neuronal defects can act as a primary trigger of glial activation. Future studies can now clarify the mechanisms of the trans-cellular interactions and their ultimate consequence on worsening neuronal function, the cell type in which AD pathophysiology begins.

Finally, our findings have therapeutic implications. They provide ‘preclinical’ validation that retromer viral vectors can be a viable gene therapy approach for late-onset AD, This may also be important for other disorders like PD, for which there is evidence that primary defects in retromer-dependent endosomal trafficking act as causal drivers of disease^28–33^. Just as importantly, our findings show that the primary effect of VPS35 overexpression is on neurons that have endosomal dysfunction, and that even in those the effect is to normalize defects induced by retromer depletion. This normalization effect, together with our findings that the therapeutic dose of VPS35 overexpression is non-toxic, supports the prediction that if globally introduced to the brain, retromer gene therapy may selectively target diseased neurons.

## METHODS

### Design and preparation of the retromer constructs

DNA constructs expressing the human retromer proteins tagged with HA or GFP at c-terminal were designed and synthesized using Thermofisher’s GeneArt portal. mRNA sequence for VPS35 was acquired from the National Center for Biotechnology Information (NCBI). The constructs were then subcloned into pcDNA 3.1 (+) Hygro vectors and eventually into AAV9 transfer plasmids for viral production. The constructs contained either one of the tags (EGFP or HA) at the C-terminal. The EGFP tagged proteins contained a polyglycine (6G) linker to reduce steric hindrance. EGFP sequence was obtained from pcDNA3-EGFP plasmid (Addgene plasmid 13031).

### AAV9 production

DNA constructs were used to drive expression of transgenes such as VPS35 and GFP ^34^. The expression cassette was flanked with adeno-associated virus serotype 2 (AAV2) terminal repeats. The cytomegalovirus chicken beta-actin hybrid promoter (“CBA” or “CAG” promoter) was used. The transgene was followed by the woodchuck hepatitis virus post-transcriptional regulatory element (WPRE) and the bovine growth hormone polyadenylation signal. Empty vector control was generated by deleting the GFP sequence. Retromer transgenes were individually subcloned into the construct in lieu of the GFP: Vps35-HA and Vps35-GFP. Each of these constructs was individually packaged into a recombinant adeno-associated virus vector, AAV9, by described methods ^35^ using capsid and helper plasmid DNA from the University of Pennsylvania ^36^. The AAV9 preparations were filter-sterilized using Millex syringe filters (Millipore) at the end of the procedure and then stored frozen in aliquots. The titering method was for encapsidated vector genomes per ml by a slot-blot method against a standard curve using the Amersham ECL Direct Nucleic Acid Labeling and Detection Systems (GE Healthcare Bio-Sciences).

### Neuronal culture

Primary mouse cortical and hippocampal neuronal cultures were implemented as described previously^37^. Neurons (1.2E+06 per well) were transduced with various doses of AAV9 (5E+9 vg/well, 2E+10 vg/well & 1E+11 vg/well; Multiplicity of infection (MOI) of 4k, 17k, & 83k respectively), 7 days after plating in a 6 well plate. The culture was maintained for 3 weeks after transduction (4 weeks total). At day 28 the neurons were lysed using RIPA buffer with protease and phosphatase inhibitors.

### Co-immunoprecipitation

Co-immunoprecipitation was performed as described previously^38,39^ with some modifications. Briefly recombinant Protein G Agarose beads (ThermoFisher cat# 15920010) were washed three times with PBS, centrifuged at 1500 g (2 min) and incubated with 3.5-5 ug of primary antibody (anti-HA, ab9110 & anti-GFP, ab290) overnight with constant rocking at 4°C. Next day the antibody coated beads were washed with cold PBS (4°C) and pelleted by spinning at 1500 g for 2 min.

Cells (or tissue) were washed twice with cold PBS and homogenized in IP buffer (50mM Tris, 100mM EDTA, 150mM NaCl, pH 7.3). Protein concentration was determined using Pierce BCA protein Assay kit (ThermoFisher cat # 23225). 300-700 ug of protein was prepared in 1100uL of IP buffer. Of this 1100ul, 100ul was saved as total lysate (input) and 1000ul was incubated with antibody coated agarose beads overnight at 4°C. Next day the samples (containing beads) were centrifuged at 1500 g for 1 min at 4°C; the supernatant was collected and labeled as the unbound fraction. Beads were washed 5 times with cold PBS. After the last wash PBS was completely aspirated and proteins were eluted from the beads and samples prepared for loading onto electrophoresis gels by adding 150uL-1X sample buffer (IP buffer 100ul buffer + 35uL of 4X NuPAGE^™^ LDS Sample Buffer ThermoFisher cat# NP0007, + 15ul of 10X Reducing Agent ThermoFisher Cat# NP0009). After agitating the samples gently they were incubated at 50°C on water bath for 30 min, mixing well every 10 min. The samples were incubated at 90°C for 5 min before loading onto NuPAGE® Bis-Tris 4-12% gels.

### VPS35 nsKO mice

VPS35 floxed mice were generated at Center for Mouse Genome Modification at UConn health. Homologous recombination was performed in mouse embryonic stem cells (mES) targeting the VPS35 gene at exons 2 to 6. The recombined gene had LoxP sites before exons 3 and after exon 5. G418 and Gancyclovir selection and nested long range PCR were used to identify targeted clones. Targeted ES cells were aggregated into morula to generate chimeric mice. Next the chimeric mice were bred with ROSA26-Flpe to remove the Frt-flanked PGKneo cassette. Neuronal-selective VPS35 knockout mice were generated by crossing mice expressing loxP-flanked VPS35 (exons 3-5) (VPS35fl/fl) with mice expressing Cre recombinase under the Camk2α promoter. Camk2a-CRE mice were obtained from Jackson labs (Stock No: 005359). All experiments involving mice were approved by the Institutional Animal Care and Use Committee of Columbia University.

### Western blots

Protein from neuronal cultures and the mouse brain CA1 region were isolated as described previously^40^. Lysates from the samples were run on NuPAGE^®^ Bis-Tris 4-12% gels, transferred onto nitrocellulose membranes using iblot and were probed with antibodies. Primary antibodies targeting the following proteins were used for probing; HA-tag (ab9110, Abcam, 1:2k), Vps35 (ab57632, Abcam, 1:1k), Vps26a (ab23892, Abcam, 1:2k), VPS26b (NBP1-92575, Novus, 1:500) Vps29 (NBP1-85288, Novus, 1:500), Vps29 (sab2501105, Sigma-Aldrich, 1:500) and β-actin (ab6276, Abcam), beta III Tubulin (neurotubulin) (ab18207, Abcam, 1:2k), NeuN (ab177487, Abcam 1:1k), GluA1 (mab2263, Millipore, 1:1k), APP-CTF (ab32136, Abcam, 1:2k), GFAP (ab68428, Abcam, 1:10k), IBA1 (ab178846, Abcam, 1:500), CD11b (ab133357, Abcam, 1:500). The primary antibodies were selected based on extensive citations by previous studies, most of the antibodies used are monoclonal antibodies with minimal non-specific signal and several of them have been validated by KO testing. IRDye^®^ 800 or 680 antibodies (LI-COR) were used as secondary with dilutions of 1:10k for 800CW, 1:15k for 680RD, and 1:25k for 680LT antibodies. Western blots were scanned using the Odyssey imaging system as described previously^41^.

### Stereotactic injection of AAV9 in mouse hippocampus

All animal procedures were approved by Columbia University IACUC. AAV9 viral particles were injected into the CA1 region of the mouse brain as described previously^42^ with slight modifications. Briefly, the surgical procedure was carried out in aseptic environment. Animals were anesthetized using ketamine / xylazine. Fur was clipped and the site cleaned with betadine scrub followed by 70% ethanol. Eyes were lubricated with an application of sterile, bland ophthalmic ointment. Preemptive analgesia was induced by carprofen and marcaine injections. Mouse was placed on infra-red warming pad in the stereotaxic frame connected to an automatic injection pump. Surgeon wore proper protective equipment and sterile gloves. Using sterile scissors, skin was cut and bone exposed. A tiny burrhole was placed in the skull at the experimental coordinates calculated using landmarks (bregma and lambda). The dura mater was pierced using a sterile needle and the syringe needle lowered to the desired z coordinate.

AAV9 VPS35-HA virus was injected through 28g needle attached to a hamilton syringe at a rate of 0.2 ul/min over a period of 5-10 min (1-3 ul volume). Control AAV9 was injected on the contralateral side. Optimal dose was identified using a dose response ranging from 4E+9 to 1E+11 vector genomes. The needle was retracted 5 minutes after the end of injection. Surgical wound was closed/sutured by a combination of skin glue and sterile stitches. Post-surgery 1 ml of normal saline was injected via intraperitoneal route, to minimize dehydration. Infra-red warming pad was used during post op recovery. Animals were monitored continuously until ambulating and once daily for the first 3 days and then twice a week until day 14 post op. Sutures were removed on day 10-14 post op. Animals were survived for 1-3 months post-surgery.

### Perfusion and tissue processing

After the survival period, the animals were perfused with saline, the brains extracted and the site of injection (dorsal CA1) was microdissected out of the brain for biochemical analysis. Animals selected for fluorescent IHC analysis were perfused with normal saline followed by 4% PFA and 30% sucrose. The brains were embedded in optimal cutting temperature (OCT Tissue-tek Cat# 62550-12) and frozen at −80°C. Frozen blocks of brains in OCT were sectioned at a Leica CM3050 S cryostat set at 12 microns thickness. Sections were transferred onto Superfrost™ Plus Microscope Slides (Fisher Cat# 12-550-15) and stored at −80°C.

### Fluorescent Immunohistochemistry (IHC)

Standard staining and microscopy techniques were used, as described previously^34,43–45^ with some modifications. All washes were done with PBS (1X-PBS containing 0.01% Sodium Azide). Briefly antigen retrieval was performed using Citric Acid buffer (10mM sodium citrate, 1.9mM citric acid, pH6.0) as describe previously^46–49^. Buffer in a coplin jar was preheated in a microwave (900W, MCD790SW) before immersing the slides. The slides were then heated for 10 minutes in intervals of 3 + 2 + 3 + 2 min with monitoring/replenishing of the buffer level in between the intervals. Buffer and slides were allowed to cool down at room temperature for 30min. Slides were washed once and incubated in 0.03% H_2_O_2_ in water for 30min. After one wash sections were permeabilized using 0.01% digitonin (wt/vol) for 10 min and then blocked in 5% donkey serum in PBS (vol/vol) for 1.5 hours, followed by incubation in primary antibodies overnight at 4°C. Primary antibodies were prepared in 1% donkey serum (in PBS) for the following proteins: GFP (ab13970, Abcam, 1:200), HA-tag (ab9110, Abcam, 1:250 & ab18181, Abcam, 1:150), NeuN (ab177487, Abcam, 1:500), GluA1 (ab183797, Abcam, 1:100), GFAP (ab7260, Abcam, 1:500), IBA1, (ab178846, Abcam, 1:500). Next day after five washes, the sections were incubated for 2 hours with secondary antibody. Secondary antibodies (ThermoFisher) conjugated with alexa fluor dyes were used at a dilution of 1:250 in 1% donkey serum (in PBS). After 3 washes Hoechst 33258 nuclear stain (ThermoFisher H3569, 1:1500 for 5 min) was used to counter stain the nuclei. After another five washes coverslips were mounted onto the slides with ProLong Gold Antifade Mountant (ThermoFisher P36934). An alternate protocol was used for staining of VPS35 (ab10099, Abcam, 1:300) (Fig. 3A). Briefly, 30 μm free-floating horizontal sections were washed three times with PBS and incubated at 4°C overnight on a rotator in 1 ml of primary antibody diluted in PBS containing 0.3% Triton X-100 (vol/vol) and 5% normal donkey serum (vol/vol). After three washes with PBS-T (0.1% Triton X-100), the sections were incubated for 1 hour with secondary antibody (Donkey-anti-goat, Alexa555 at 1:400, ThermoFisher). Following three washes with PBS-T, and one wash with PBS, sections were mounted on slides. All images were captured using Zeiss LSM 700 META confocal microscope equipped with a 5x, 20x & 63× Plan-Apochromat objective and HeNE1, HeNe2 and argon lasers.

### Statistics

Statistical analysis was performed using Microsoft Excel and SPSS. Student’s t-test was used for all biochemical experiments assuming equal variance. Independent two-sample t-test with two-tailed distribution and paired t-test were used depending on the type of experimental data. All data are presented as means, the error bars indicate standard error of the mean. All bar graphs were created in GraphPad Prism 8.

## Acknowledgements

This study was partly supported by an NIH grant AG008702 and an anonymous foundation to S.A.S. and sponsored research from Meira GTX to G.A.P. This paper is dedicated to the memory of Ronald L. Klein

## Supplementary Figure

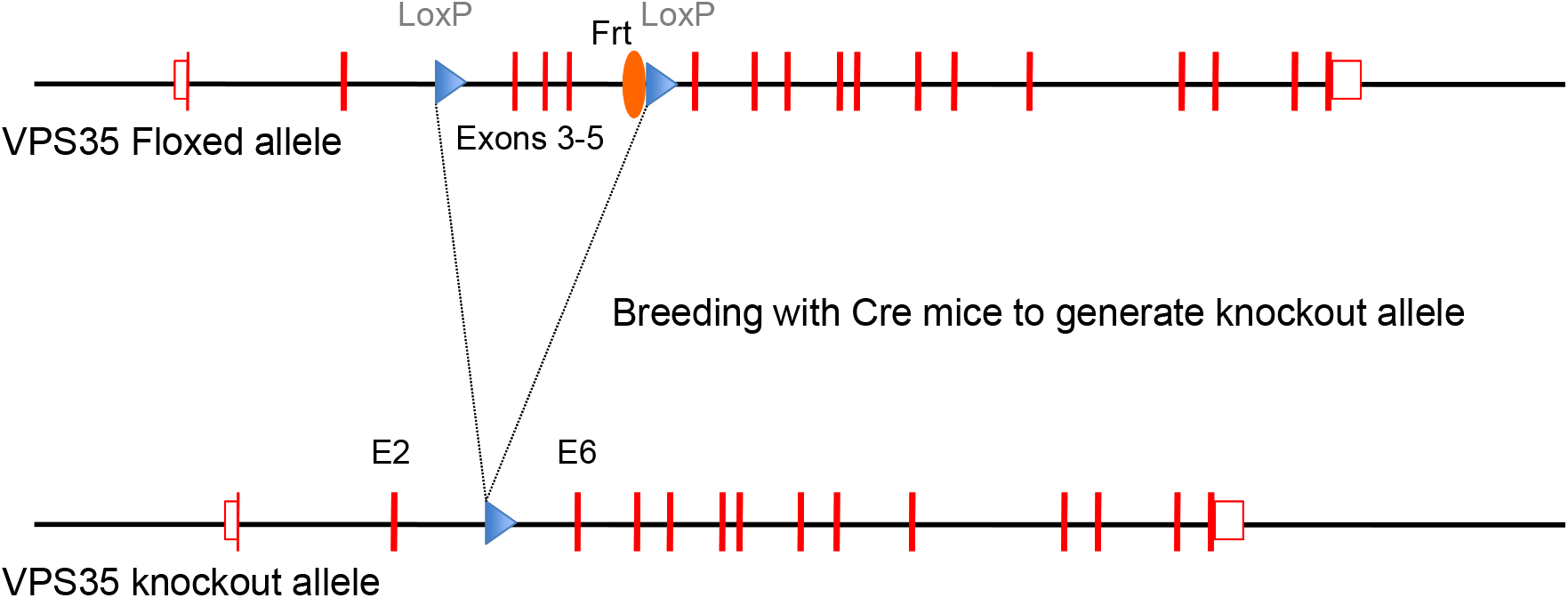
Homologous recombination was performed in mouse embryonic stem cells (mES) targeting the VPS35 gene at exons 2 to 6. The recombined gene had LoxP sites before exons 3 and after exon 5. G418 and Gancyclovir selection and nested long range PCR were used to identify targeted clones. Targeted ES cells were aggregated into morula to generate chimeric mice. Next the chimeric mice were bred with ROSA26-Flpe to remove the Frt-flanked PGKneo cassette. Neuronal-selective VPS35 knockout mice were generated by crossing mice expressing loxP-flanked VPS35 (exons 3-5) (VPS35fl/fl) with mice expressing Cre recombinase under the Camk2α promoter. Camk2a-CRE mice were obtained from Jackson labs (Stock No: 005359)

